# HierCC: A multi-level clustering scheme for population assignments based on core genome MLST

**DOI:** 10.1101/2020.11.25.397539

**Authors:** Zhemin Zhou, Jane Charlesworth, Mark Achtman

## Abstract

**Motivation:** Routine infectious disease surveillance is increasingly based on large-scale whole genome sequencing databases. Real-time surveillance would benefit from immediate assignments of each genome assembly to hierarchical population structures. Here we present HierCC, a scalable clustering scheme based on core genome multi-locus typing that allows incremental, static, multi-level cluster assignments of genomes. We also present HCCeval, which identifies optimal thresholds for assigning genomes to cohesive HierCC clusters. HierCC was implemented in EnteroBase in 2018, and has since genotyped >400,000 genomes from *Salmonella, Escherichia, Yersinia* and *Clostridioides*.

**Availability:** Implementation: http://enterobase.warwick.ac.uk/ and Source codes: https://github.com/zheminzhou/HierCC

**Supplementary information:** Supplementary data are available at *Bioinformatics* online.

## 1 Introduction

Following its introduction in 2011 (Mellmann et al., 2011), core genome multi-locus sequence typing (cgMLST) was widely adopted as a scalable, portable and easily communicable genotyping solution for the genome-based, routine surveillance of bacterial pathogens (Jolley et al., 2012; Jones et al., 2019; Moura et al., 2016). In a cgMLST scheme, bacterial genomes are assigned to sequence types (STs) consisting of 1000s of integers, which each represents a distinct sequence variant (allele) of a soft core gene (Zhou et al., 2020). A pairwise comparison of allelic differences between STs approximates the genetic distance between genomes, and can be used for downstream phylogenetic analyses. However, STs are arbitrary constructs, and natural bacterial populations can each encompass multiple, related STs.

Several single-level clustering schemes have been applied to cgMLST schemes to extract single-level clusters (SCs) from a hierarchical clustering. Such SCs were equated with sub-lineages in *Listeria* (Moura et al., 2016) or single source outbreaks of *Salmonella* serovar Enteritidis (Coipan et al., 2020). However, because SC schemes identify only one clustering level, they ignore the wide spectrum of genetic diversities and the existence of multiple hierarchical levels of natural populations.

Unlike MLST schemes, multiple multi-level clustering (MC) schemes for bacterial pathogens exist that are based on core genomic SNPs. For example, SnapperDB assigns *Salmonella* genomes to so-called SNP addresses, consisting of seven hierarchical single-linkage clusters based on SNP distances (Dallman et al., 2018). Similarly, genomes of *Yersinia pestis*, or *Salmonella* Typhi are assigned to one of multiple levels of sublineages based on their placement in a phylogeny (Morelli et al., 2010; Wong et al., 2016). However, SNP-based approaches are restricted to relatively uniform clades because, unlike cgMLST schemes which can extend to the genus level, the SNP calls and phylogenetic reconstructions become less reliable at higher levels of intra-genus diversity.

Here we present HierCC, a scalable scheme that assigns bacterial genomes in real-time to multi-level clusters spanning a wide spectrum of genetic diversities. We also present HCCeval, which identifies optimal levels from the HierCC results, and yields multi-level clusters that likely represent hierarchical natural bacterial populations up to the species level.

## 2 HierCC Workflow

HierCC firstly calculates a minimum spanning tree (MST) (Fig. 1) based on a special distance metric that minimizes the topological distortion due to missing genes (Supplementary Text). The resulted tree is then used to assign every genome into multi-level clusters. Alternatively, HierCC runs a so called “production mode”, in which case every new genome is attached into a pre-calculated tree without changing the cluster assignment of any existing genome (Supplementary Text).

**Fig. 1.**
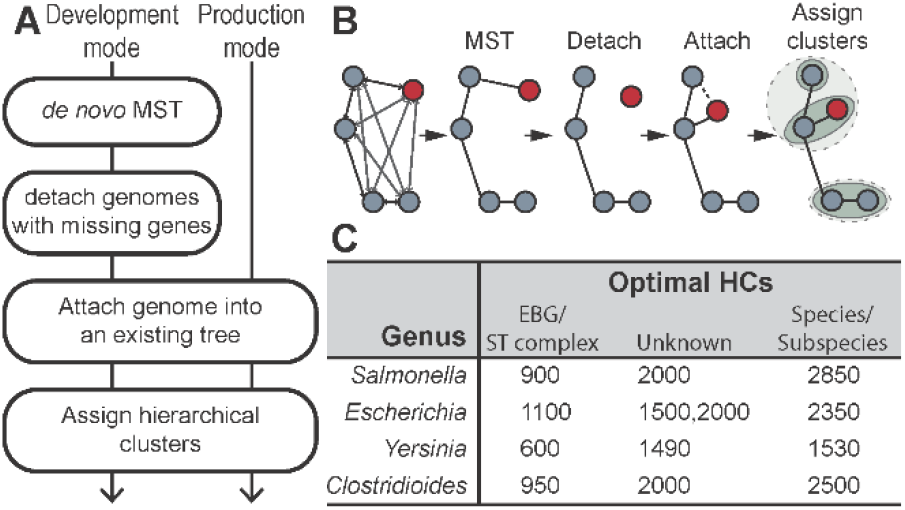
(**A**) The workflows of HierCC in complete or production mode. (**B**) Cartoon of the workflow in complete mode. The node in red indicates a genome that carries numerous missing genes. (**C**) The optimal HC levels identified by HCCeval in 4 EnteroBase databases.

## 3 Evaluation of HCs using HCCeval

Some of the HC levels generated by HierCC may correspond to genetically isolated, natural populations. HCCeval was designed to identify those HC levels that best reconcile natural distinct populations. Given a set of STs and their corresponding HierCC results, HCCeval tests the stability of every HC level by comparing its clusters with the clusters from all other levels using the normalized mutual information (NMI) score. Visual inspection of a heat map of NMI scores allows the identification of “stable blocks”, which consist of sets of continuous HC levels that define highly similar clusters (NMI >0.9). For every HC level, HCCeval also calculates a silhouette score, which evaluates the cluster cohesiveness by comparing the internal pairwise similarities of each genome in a cluster with its similarities to genomes from other clusters. The clusters in HC levels with the greatest silhouette score for each stable NMI block are likely to represent natural populations (Fig. S1).

## 4 Implementation in EnteroBase

An initial set of hierarchical clusters was calculated in 2018 by EnteroBase HierCC in development mode for representative genomes from all its genome databases with a cgMLST scheme (*Salmonella, Escherichia, Yersinia* and *Clostridioides*). These representatives consisted of one genome per ribosomal MLST type, and their hierarchical clusters were evaluated by HCCeval (Fig. S1 and (Frentrup et al., 2019)). Visual inspection of the results identified 3-4 stable blocks for each genus. The highest HierCC clusters (HC1530-HC2850 depending on genus) correspond to subspecies or species (Figure 1C) (http://enterobase.readthedocs.io/en/latest/HierCC_lookup.html). The lowest clusters (HC600-HC1100) are consistent with eBGs or ST complexes or comparable populations defined by 7-gene MLST schemes (Frentrup et al., 2019; Zhou et al., 2020). Additionally, EnteroBase also publishes all genome cluster assignments for HC0, HC2, HC5, HC10, HC20, HC50, HC100, HC200 and HC400. Subsequent experience has indicated that some these HC levels correlate with infectious outbreaks (Jones et al., 2019) or recent local transmissions (Frentrup et al., 2019; Zhou et al., 2020).

## 5 Conclusions

In this article, we introduce HierCC, a scalable, fine-grained, incremental clustering scheme for bacterial genomes based on their cgMLST allelic profiles. HierCC was integrated into EnteroBase in 2018,and has currently assigned >400,000 genomes from *Salmonella, Escherichia, Yersinia* and *Clostridioides* into 12-13 multi-level clusters from sub-clonal variation to species. HierCC is offered as a stand-alone package that is suitable for any bacterial genus with a cgMLST scheme where large numbers of genomes are available for the initial assignments.

## Supporting information

Figure S1. Statistical evaluation of HierCC at all levels for EnteroBase databases of Salmonella (A), Escherichia (B) and Yersinia (C).

Supplementary Text: the workflow of HierCC

## Acknowledgements

We gratefully acknowledge feedback for *Salmonella* HierCC schemes from Francois-Xavier Weill and María Pardos de la Gándara, Institut Pasteur, and the maintenance of EnteroBase by Khaled Mohamed.

## Funding

This project was supported by the Wellcome Trust (202792/Z/16/Z) and the BBSRC (BB/L020319/1).

## Conflict of Interest

none declared.

## Notes

https://enterobase.warwick.ac.uk

### Competing Interest Statement

The authors have declared no competing interest.

