## Supplementary material for "HierCC: A multi-level clustering scheme for population assignments based on core genome MLST": Figure S1. Statistical evaluation of HierCC at all levels for EnteroBase databases of Salmonella (A), Escherichia (B) and Yersinia (C).

A. *Salmonella* HierCC

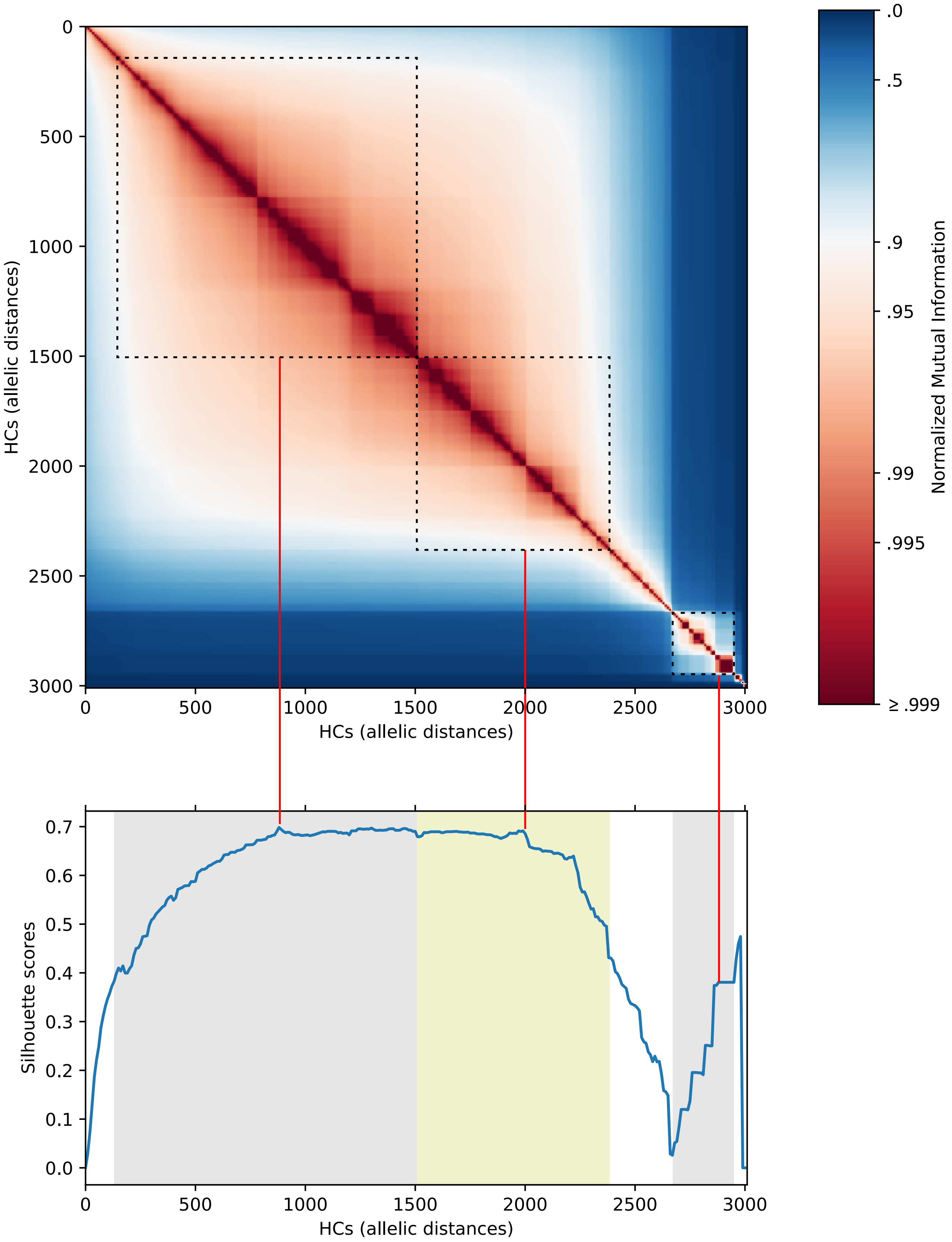


Supplementary Figure 1. Statistical evaluation of HierCC at all levels for EnteroBase databases of *Salmonella* (A), *Escherichia* (B) and *Yersinia* (C). The heatplot at the top of every figure shows Normalized mutual information score from pairwise comparisons of clusters at different hierarchical levels, and the line plot at the bottom shows the silhouette score of the clusters at every hierarchical level. The dotted boxes and the shaded area indicates ranges of stable clustering, and the red lines indicate chosen hierarchical levels of maximum cluster stability for each HierCC scheme.

B. *Escherichia* HierCC

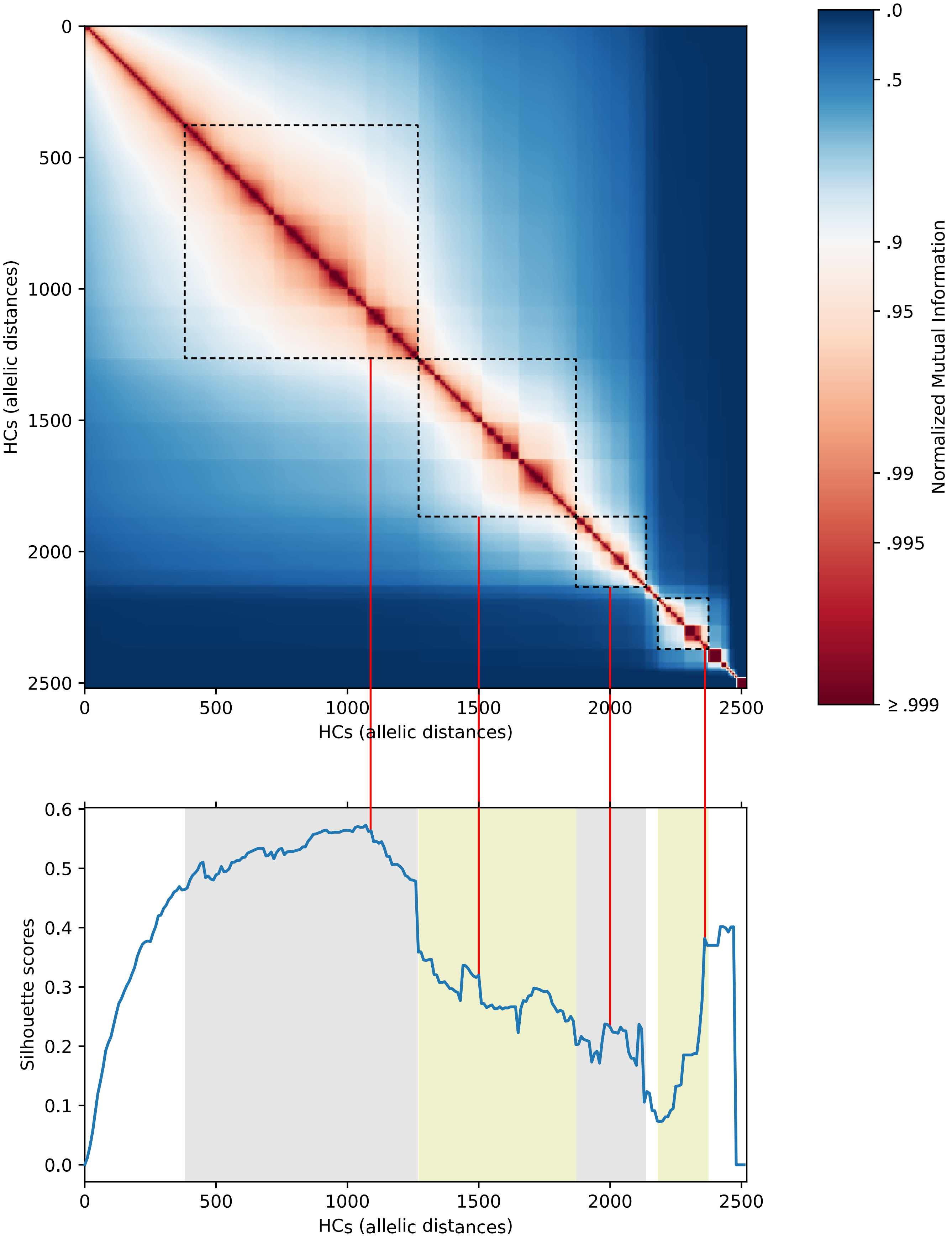


Supplementary Figure 1. Statistical evaluation of HierCC at all levels for EnteroBase databases of *Salmonella* (A), *Escherichia* (B) and *Yersinia* (C). The heatplot at the top of every figure shows Normalized mutual information score from pairwise comparisons of clusters at different hierarchical levels, and the line plot at the bottom shows the silhouette score of the clusters at every hierarchical level. The dotted boxes and the shaded area indicates ranges of stable clustering, and the red lines indicate chosen hierarchical levels of maximum cluster stability for each HierCC scheme.

C. *Yersinia* HierCC

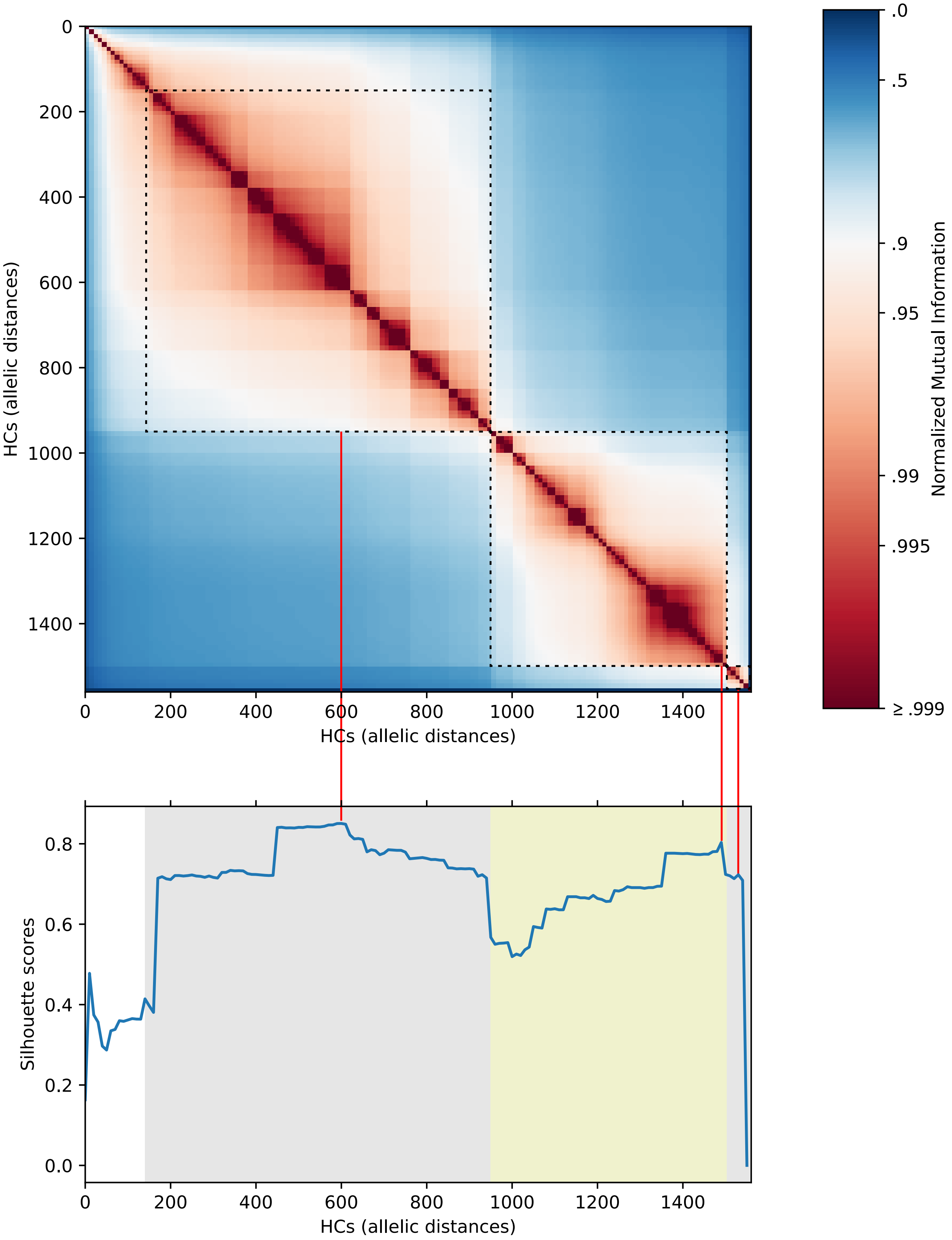


Supplementary Figure 1. Statistical evaluation of HierCC at all levels for EnteroBase databases of *Salmonella* (A), *Escherichia* (B) and *Yersinia* (C). The heatplot at the top of every figure shows Normalized mutual information score from pairwise comparisons of clusters at different hierarchical levels, and the line plot at the bottom shows the silhouette score of the clusters at every hierarchical level. The dotted boxes and the shaded area indicates ranges of stable clustering, and the red lines indicate chosen hierarchical levels of maximum cluster stability for each HierCC scheme.
