## Supplementary Text: the workflow of HierCC for "HierCC: A multi-level clustering scheme for population assignments based on core genome MLST"

### 1 *de novo* MST construction (Development mode only)

HierCC uses the Python SciPy package to generate a minimum spanning tree (MST) from a distance matrix of the numbers of differing alleles in pairwise comparisons (Fig. 1). However, excessive missing genes can artificially distort a distance matrix toward conserved, smaller genes because larger genes provide more opportunities for deletion. We arbitrarily chose to deal separately with that tiny subset of all pairs of genomes (<1%) which differ by more than 3% of missing data in order to minimize these distorting effects. Given a cgMLST scheme with *n* core genes, such problematical pairs were identified by calculating s, a minimum number of shared genes between pairs of genomes as:

$$s=m-0.03*n$$

where $m$ is the number of core genes in the more complete genome. The allelic distance *d* between two genomes is calculated as:

$d=\left\{ \begin{aligned} \left\lfloor n*{d_{0}}/{s_{0}}+0.5 \right\rfloor, s_{0}\geq s \\ \left\lfloor n*{(d_{0}+s-s_{0})}/s+0.5 \right\rfloor, s_{0}<s \end{aligned} \right.$	(1)

where *s_0_* is the number of core genes common to both genomes and *d_0_* is the number of different alleles. In the second case, HierCC elevates the allelic distances between the two genomes, thereby increasing the likelihood that genomes with more missing data remain outliers and do not join tight clusters.

After completion of the MST, HierCC actively removes genomes with >3% of missing genes from the MST, and deals with them in the next step.

### 2 Attach nodes to MST

Given a detached genome *q* and a genome *r* in the MST, HierCC calculates a directional distance from *r* to *q* as:

$d_{r\to q}^{'}=\left\{ \begin{aligned} \left\lfloor n*{d_{0}}/{s_{0}}+0.5 \right\rfloor, s_{0}\geq s' \\ \left\lfloor n*{(d_{0}+s'-s_{0})}/{s'}+0.5 \right\rfloor, s_{0}<s' \end{aligned} \right.$	(2)

Equation 2 only differs from equation 1 by its use of *s’*, which is calculated as:

$$s’=m_{q}-0.03*n$$

where *m_q_* is the number of core genes in *q*. Equation 2 may give a lower distance measure than equation 1 when there are more missing genes in *q* than in *r*. An edge is then drawn between *q* and a genome *r* in the tree that minimizes $d_{r\to q}^{'}$, or to the genome with the smallest (oldest) ST designation when multiple STs are equally distant.

### 3 Assign each genome to hierarchical clusters

HierCC applies every threshold $c\in\{0..n\}$ to a copy of the MST by removing all edges with greater distances than the threshold. All remaining, discrete subtrees constitute the clusters for that threshold, which are designated by the smallest ST designation in the cluster. As a result, each genome is assigned to *n +1* clusters, from Hierarchical Cutoff 0 (HC0) which groups together genomes that have no allelic difference except for missing data, to HC*n* which consists a single cluster of all genomes.

### 4 Production mode

When the HierCC pipeline loads an optional existing tree together with additional new genomes, HierCC changes to “production mode”, and only performs the workflows in sections 2,2 and 2.3 without recalculation of an MST (Fig. 1A). In production mode, cluster assignments of previously assigned genomes are not modified because of subsequent attachment of new genomes to the tree. One consequence of this behavior, is that the gradual merging of clusters is prohibited which is so typical of ST complexes in SC assignments of 7-gene MLSTs. However, we note that the resulting tree is no longer an MST, and genetically closely related genomes may be assigned to separate clusters.
